# Structure of Lekking Courts of Male White-Bearded Manakins (*Manacus Manacus*) Is Linked to Their Attractiveness

**DOI:** 10.1101/226944

**Authors:** Kirill Tokarev, Yntze van der Hoek

**Affiliations:** Dept. of Psychology, Hunter College, City University of New York, New York, NY, USA; Dept. of Radiology, Weill Cornell Medicine, New York, NY, USA; Universidad Regional Amazónica IKIAM, Tena, Ecuador

## Abstract

Male White-bearded Manakins (*Manacus manacus*) perform courtship displays on individual courts in close proximity of each other, while females visit them to choose potential mates. These displays represent a sequence of physiologically demanding movements, including rapid hops between saplings on the court. Our observations of courtship behavior and court characteristics of eight males suggest that the structure of the court may be an important factor in courtship: we found regularities in the inter-sapling distances on the courts of males that attracted females. We hypothesize that sexual selection by females may favor those males that have courts providing an optimal platform for their courtship display.

**Estructura de escenarios de lek de saltarines barbiblancos (Manacus manacus) es conectada a su atractivo:** Resumen: Los saltarines barbiblancos (Manacus manacus) machos realizan despliegues de cortejo en “escenarios” individuales cercanos los unos a los otros, mientras las hembras los visitan para escoger su pareja potencial. Estos despliegues representan una secuencia de movimientos fisiológicamente arduos, que incluyen saltos rápidos entre los palos o plantones del escenario. Nuestras observaciones de comportamiento nupcial y de las características de los escenarios de ocho machos sugieren que la estructura del escenario puede ser importante para cortejo: habían regularidades en distancias entre los palos de los escenarios de los machos que atrajeron hembras. Formulamos la hipótesis que la selección sexual realizada por las hembras favorece los machos con escenarios que proporcionan una plataforma óptima para sus despliegues de cortejo.

## INTRODUCTION

Male White-bearded Manakins (*Manacus manacus*) engage in courtship displays in leks, with males on individual courts but within visual and auditory range of the neighbors (Darnton 1958; Snow 1962; Thery 1992). Such aggregations are both cooperative and competitive, as several males lekking together are more likely to attract females, but only very few of these males gain the vast majority of matings (Lill 1974; Shorey et al. 2000; Shorey 2002). Previous work on lekking birds, and manakins in particular, has identified various morphological, behavioral and territorial factors important for male mating success (Fiske et al. 1998; Fusani et al. 2014; Shorey 2002). Male courtship display of bearded species of manakins is very physiologically demanding, as it involves a series of rapid hops between the saplings on the court, when males use only the power of legs for movement while wings are used to produce a snapping sound (Barske et al. 2011; Barske et al. 2014; Fusani et al. 2014); males speed up their performance in presence of females (Barske et al. 2015). Thus, such male courtship displays may represent an honest signal of males’ motor skills for sexual selection by females. Manakin males that attract more females tend to have centrally located courts within the lek (Shorey 2002; Stein & Uy 2006). Although male manakins clear the floor of their courts, potentially to increase visual appeal of their display (Uy & Endler 2004; Chiver & Schlinger 2017) and as anti-predation strategy (Cestari & Pizo 2014), and have a favorite sapling for mating (Coccon et al. 2012), structure of their courts has not been studied in relation to mating success. In the present work, we provide observations suggesting that attractive males tend to have courts with a specific arrangement of saplings on them.

## METHODS

From February to April 2017, we studied lekking behavior of White-bearded Manakins on private land governed by the Kichwa community Tamia Yura in Napo Province, Ecuador (0°58’S, 77°47’W; 650 m a.s.l.; **Figure 1**). A lek consisting of at least eight males was located in secondary lowland tropical moist forest on the fringe of mixed agroforestry land, an area with dense understory vegetation comprised of shrubs and trees belonging to a large variety of angiosperm families such as *Fabaceae, Lauraceae, Rubiaceae,* and *Melastomataceae.*

**Figure 1.**
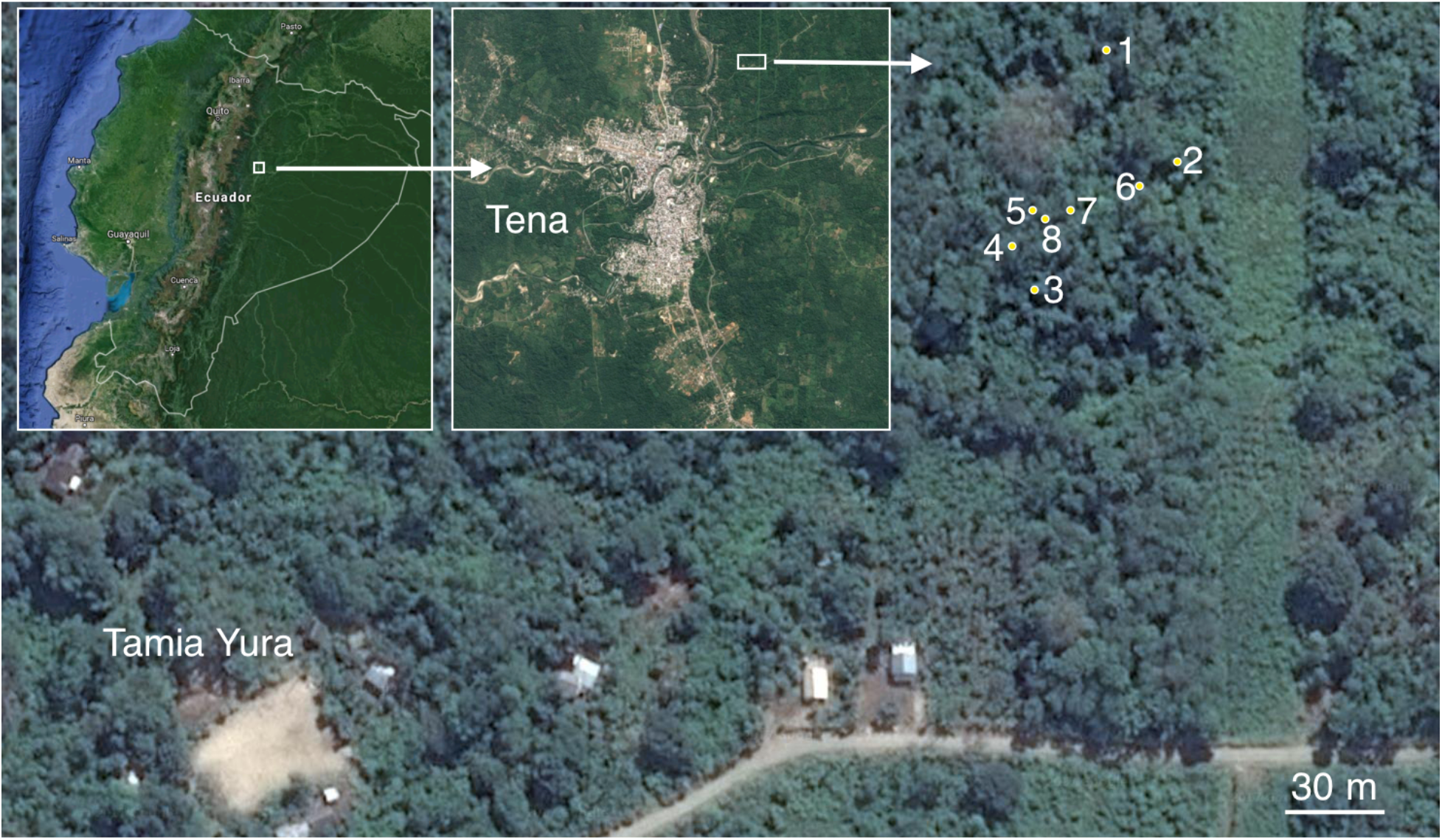
The study area and locations of individual display courts within the studied lek of White-bearded Manakins. The courts are marked from 1-8, with court 1 being used by a juvenile; female visits were observed only at courts 6, 7 and 8. The inserts show the location on the map of Ecuador and the vicinity of the city of Tena. The image was produced using ‘Google Maps’.

We observed the behavior of adult and juvenile males and females at the lek between 06:00 and 18:00 (ECT) for a total of 140 h, over the course of 21 days; video recording of 15 h. Two ‘GoPro Hero4’ cameras were used at a time; they were set remotely through an iPhone app in order to minimize our movement at the lek, a possible influence on the birds’ behavior. Vocalizations and wing snaps made during courtship display allowed us to follow which males were active, reducing potential biases in missing court visits by females. However, it should be noted that we did not observe or record activity at the courts continuously 24h every day, and thus our data might reflect the probability of female visits rather than absolute qualities of mating success (which ideally would include identification of offspring). Female visit was counted when a female perched on one of the saplings that male used in his courtship displays.

Measurements of the distances between saplings used for lekking display were taken for each court in the end of the observation period. From these measurements, we calculated the mean distance and used nonparametric Mann–Whitney U-test to assess whether inter-sapling distance at courts visited by females differed from distances found at unvisited courts. We used Kernel distribution plots to evaluate the relationship between inter-sappling distances and attractiveness of male courts (using ggplot2 package in R <u>https://CRAN.R-project.org/package=ggplot2</u>).

In addition to that, we also did measurements of the whole lek, as well as the distance to a neighboring lek that we found in the end of our observations. Each court was photographed with iPhone 5s, and GPS coordinates were extracted from the metadata of the photographs to produce the map over a satellite image online using ‘Google Maps’. The online tools of ‘Google Maps’ were also used to measure distances between the courts and the total area of the lek, as well as the distance to another lek (we do not have more data for it, as we only found it in the end of our observations). The polygonal area enclosed by male display courts was used as the minimum-lek-area (MLA) to estimate the lek size (Olson & Mcdowell 1983).

## RESULTS

The courts of the Tamia Yura lek observed in this work were delimited by 3-4 saplings. Distances between these saplings varied from 35 to 148 cm (**Figure 2**). Per court, we found that mean inter-sapling distances varied between 58 and 119 cm, with the largest mean distance found in the only court held by a juvenile male (court 1). Among the adult males, the inter-sapling distance was approximately 71±9 cm (mean ± SE). We observed that the courts visited by females contained two to three inter-sapling distances within that mean range (courts 6, 7 and 8, datapoints between two horizontal grey lines in **Figure. 2**), whereas other courts had zero (juvenile, court 1), one (courts 2, 3 and 4) or two (court 5) distances within the mean range. In addition, only the courts visited by females had at least two inter-sapling distances that were almost equal (difference ≤3 cm, overlapping circles in **Figure. 2**).

**Figure 2.**
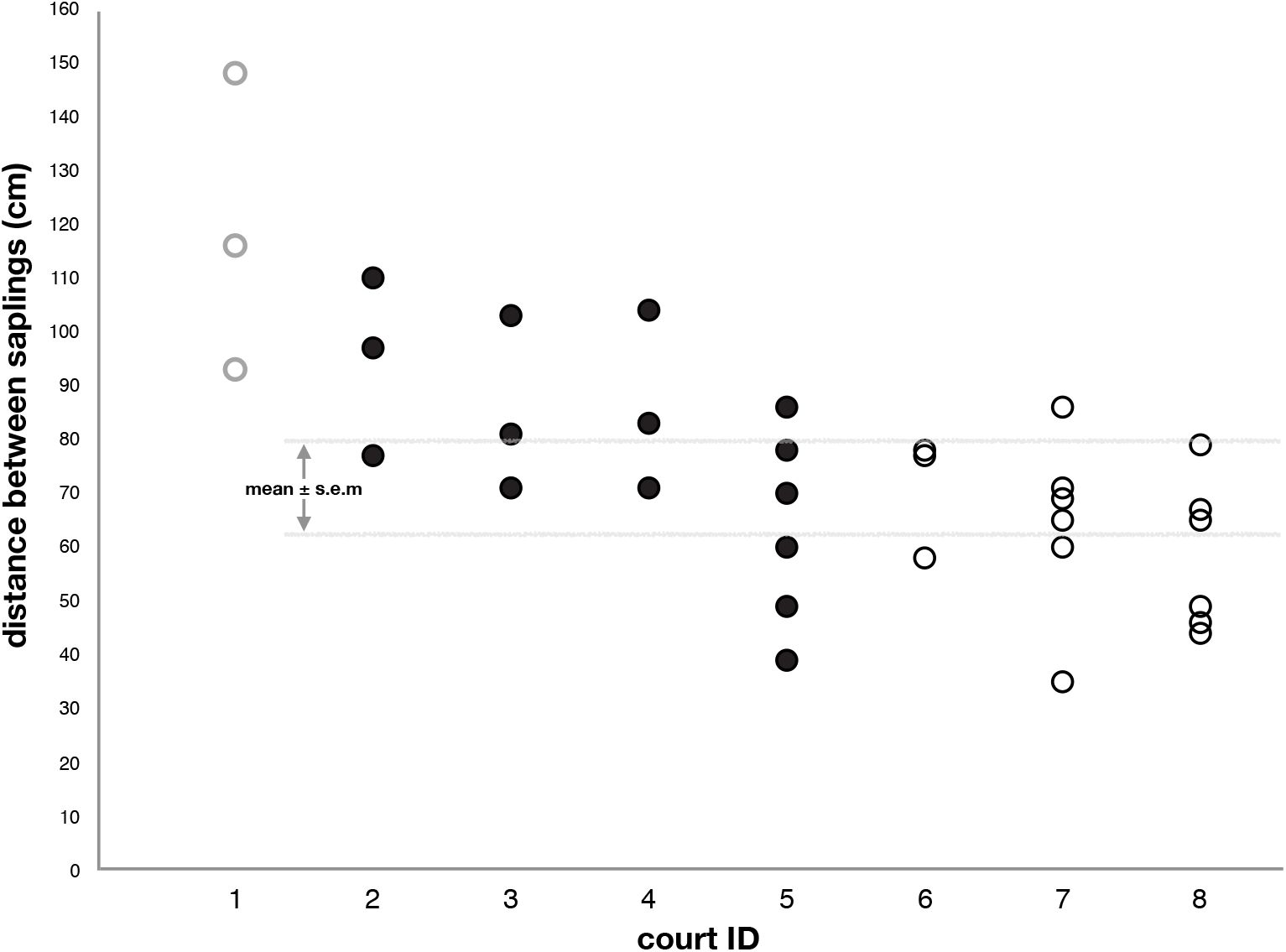
Distances between saplings in individual lekking courts. Data from each court is shown (marked from 1-8): grey open circles show values from the only juvenile court found; black filled circles – courts of adult males not visited by females; black open circles – courts of adult males that were visited by females. Horizontal grey lines represent the mean ± standard error of inter-sapling distances, calculated for adult courts.

The distribution of inter-sapling distances in courts visited by females was more uniform than distribution of inter-sapling distances at unvisited courts (**Figure. 3**). Mean inter-sapling distance at courts visited by females was significantly shorter (63 cm, CI 95% [72.6, 98.2]) than that found at unvisited courts (85 cm, CI 95% [55.17, 71.36])(p = 0.004, Mann–Whitney U test).

**Figure 3.**
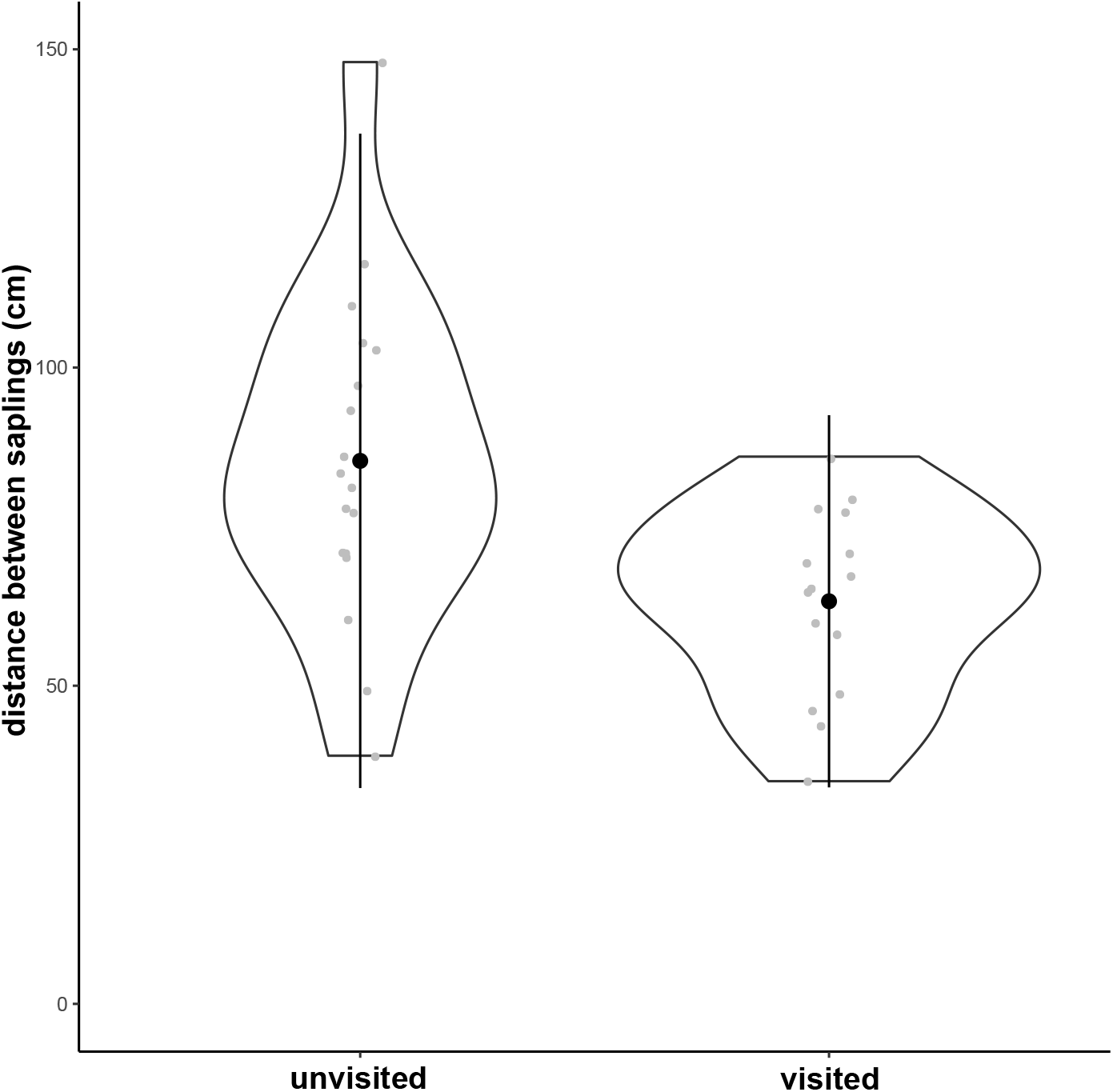
Comparison of the distributions of inter-sapling distances in the lekking courts in relation to female visits. Violin plots show kernel density of the distribution of inter-sapling distances in the lekking courts that were either unvisited *(left)* or visited by females *(right).* Black circles and vertical lines show mean ± SD (significantly different; p = 0.004, Mann–Whitney U test). Gray circles show individual datapoints.

The area delineated by the lekking courts (minimum-lek-area) was 1553 m^2^. The mean distance between courts was 35 ± 19 m; the only court held by a juvenile male was relatively isolated from the rest (**Figure. 1**). Females were only observed at courts 6, 7 and 8, which were more centrally located compared to more peripheral courts 1-5 (**Figure. 1**). Lek characteristics in comparison to other leks known within the species range are presented in **Table 1**.

**Table 1.**
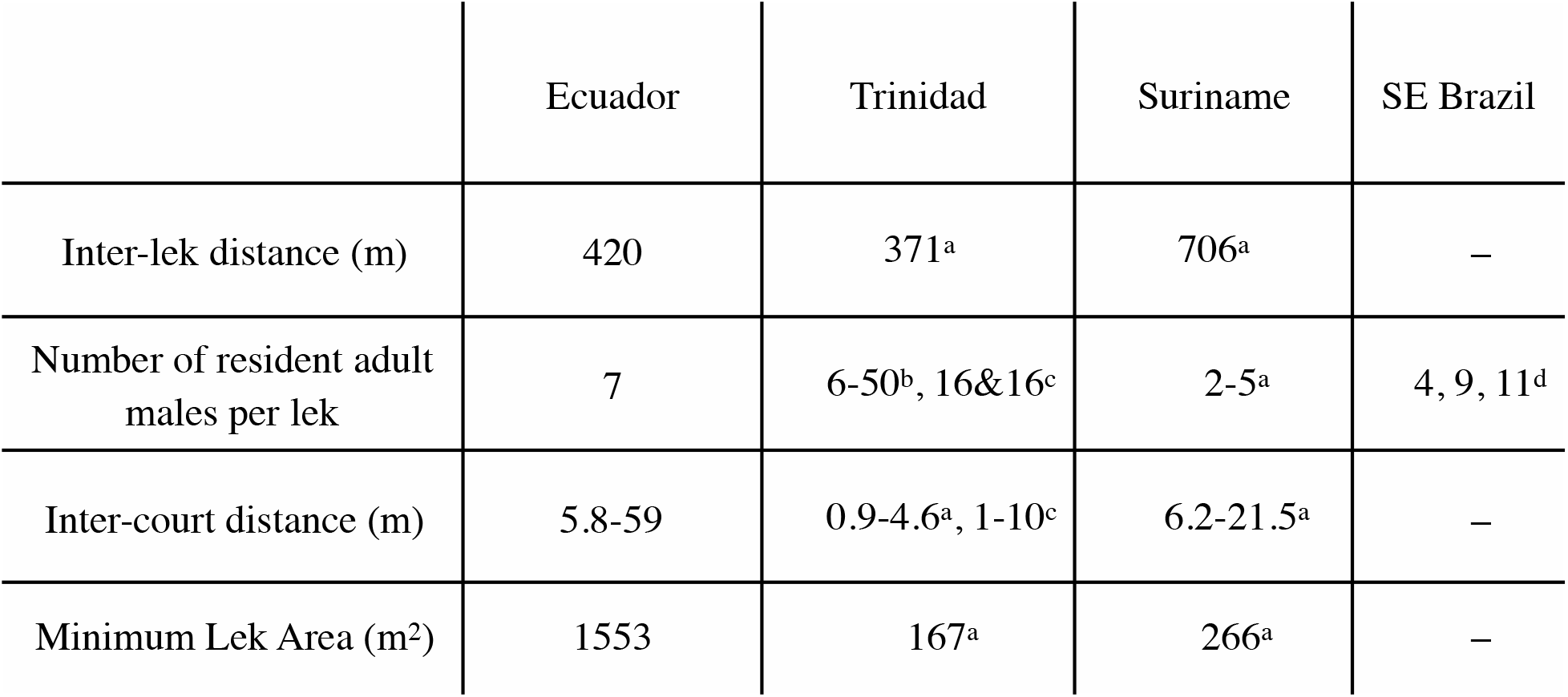
Characteristics of White-bearded Manakin leks across the species range in the Neotropics. a) (Olson & Mcdowell 1983); b) (Lill 1974); c) (Shorey 2002); d) (Thery 1992)

|  | Ecuador | Trinidad | Suriname | SE Brazil |
| --- | --- | --- | --- | --- |
| Inter-lek distance (m) | 420 | 371<sup>a</sup> | 706<sup>a</sup> | – |
| Number of resident adult males per lek | 7 | 6-50<sup>b</sup>, 16&16<sup>c</sup> | 2-5<sup>a</sup> | 4, 9, 11<sup>d</sup> |
| Inter-court distance (m) | 5.8-59 | 0.9-4.6<sup>a</sup>, 1-10<sup>c</sup> | 6.2-21.5<sup>a</sup> | – |
| Minimum Lek Area (m<sup>2</sup>) | 1553 | 167<sup>a</sup> | 266<sup>a</sup> | – |

## DISCUSSION

Our observations suggest that the structure of the lekking courts of White-bearded Manakins may be an important factor in courtship success. Phenotypic and behavioral characteristics of male manakins and the location of their courts within leks are linked to mating success (Fusani et al. 2014; Shorey 2002; Thery 1992). Centrality of the position of a court within a lek increases mating success in White-bearded (Shorey 2002) and Golden-collared Manakins (Stein & Uy 2006), as well as other avian and mammal lekking species (Fiske et al. 1998; Höglund & Robertson 1990; Kruijt & Hogan 1967). Our observations are in line with these studies, as we also observed female visits in centrally located courts but not in peripheral ones. However, we also found other regularities in characteristics of the courts that attracted females. The courts that were visited by females tended to have more saplings at a distance of 71±9 cm; also, at least two distances between saplings in such courts were of almost equal length. Likely, deviation from this mean distance, or more variation in distances between saplings at a court, leads to decreased display performance.

Such court characteristics as area and number of saplings do not influence mating success in the White-bearded Manakin (Shorey 2002). Manakins prefer saplings of ≤3m in height for perching (Mallorquin & Quevedo 2002), and although secondary tropical forest provides plenty of such saplings (Snow 1962), it seems important that the courts would have them at a certain distance from each other, because the males must be able to hop between them at speeds limited by their physiological capacities (Barske et al. 2011; Barske et al. 2014; Barske et al. 2015; Fusani et al. 2014). Our data suggest that there may be stabilizing selection towards a standard, optimal distance between saplings for such acrobatic displays.

Also, it appears that male manakins practice their performance (Coccon et al. 2012), but females may challenge them while visiting the courts (Barske et al. 2015; Fusani et al. 2014). It may be easier for a male with a possibility to do similar jumps between more than one pair of saplings. Indeed, we found that at least two pairs of saplings were almost equidistant in the courts that attracted females but not in the unvisited courts.

Thus, we suggest that attractive males not only select (and potentially protect) centrally located courts (Höglund & Robertson 1990; Shorey 2002; Stein & Uy 2006), but also choose them according to their geometric structure, which would enable the demanding courtship display at the optimal level. Future studies should compare characteristics of male courtship display, e.g. its rhythm and velocity, with court structure directly.

## ACKNOWLEDGMENTS

We thank Tamia Yura (<u>https://tamiayura.wordpress.com</u>) community, particularly Melida Tapuy, Benjamin Mamallacta Almarado, Ladi Tapuy and Carlos Cerda, for their help in finding the leks and facilitation of the logistics. We acknowledge Ecuador’s Ministry of Environment (MAE) for granting us research permit 06-16-IC-FAU-DPAN/ MA to conduct fieldwork near Tena, Ecuador.

